# Another step towards understanding evolutionary changes in *Bruggmanniella* Tavares, 1909 group (Diptera, Cecidomyiidae, Asphondyliini)

**DOI:** 10.1101/2021.09.10.459830

**Authors:** Carolina de Almeida Garcia, Carlos José Einicker Lamas, Maria Virginia Urso-Guimarães

**Affiliations:** Laboratório de Diptera, Museu de Zoologia da Universidade de São Paulo, Av. Nazaré, 481, Ipiranga, São Paulo, SP; Laboratório de Sistemática de Diptera, Departamento de Biologia, Universidade Federal de São Carlos, Rodovia João Leme dos Santos, Km 110, Bairro do Itinga, Sorocaba, São Paulo

## Abstract

An update of the delimitation of the genus *Bruggmanniella* based on phylogenetic analysis using morphological data is presented. We included the seven new species of *Bruggmanniella* described between 2019 and 2020, and discuss some aspects of the evolutionary changes among the closely related genera *Bruggmanniella, Pseudasphondylia*, and *Illiciomyia. Bruggmanniella* is confirmed here as a monophyletic Neotropical lineage, divergent from the Asian species. The phylogenetic reconstruction hypothesized here reinforces the pertinence of the genus *Odontokeros* to house all species occurring in the Oriental/Palearctic region under *Bruggmanniella*. The delimitation of *Bruggmanniella*, the geographical distribution, and niche occupation are discussed.

## Introduction

Garcia et al. [1] proposed a phylogenetic hypothesis for *Bruggmanniella* Tavares, 1909 based on morphological data of all life stages of twelve species described to date. To understand the evolution of the genus, the outgroup included six species of the Oriental genus *Pseudasphondylia* Monzen, 1955 and the species of the monotypic genus *Illiciomyia* Tokuda, 2004. The three genera share two separate teeth of the gonostylus and transverse rows of strong spines on the anterior half of the pupal tergites, indicating a close relationship among them, widely reported in the literature [2–6]. The analysis of Garcia et al. [1] supported the transference of the Oriental species of *Bruggmanniella* to *Pseudasphondylia* and the erection of a new genus, *Odontokeros* Garcia et al., 2020 to house *B. brevipes* Lin, Yang & Tokuda, 2019, the only known species of the group recorded from Taiwan at time. According to these results, *Bruggmanniella* is an endemic genus of the neotropics, while *Pseudasphondylia* and *Illiciomyia* would be restricted to the Oriental/Palearctic regions.

Simultaneously, seven new species of *Bruggmanniella* were described, *B. litseae*, *B. sanlianensis, B. shianguei*, and *B. turoguei* from Taiwan [7, 8], and *B. miconiae, B. notatae*, and *B. sideroxyli* from Brazil [9]. Lin et al. [8] also proposed an hypothesis for *Bruggmanniella* based on a molecular analysis allegedly conflicting with the hypothesis of Garcia et al. [1].

In this paper, we included the seven new species in *Bruggmanniella* and discuss some aspects of the evolutionary changes in the group of Asphondyliina with two-toothed gonostyli, which includes *Bruggmanniella*, *Pseudasphondylia*, and *Illiciomyia*.

## Material and Methods

### Taxa sampling and phylogenetic analysis

The phylogenetic hypothesis presented next is mainly derived from Garcia et al. [1]. To update the analysis, we included the recently described species of *Bruggmanniella: B. miconiae* Rodrigues & Maia; *B. notatae* Rodrigues & Maia; *B. sideroxyli* Rodrigues & Maia; *B. litseae* Lin, Tokuda & Yang; *B. sanlianensis* Lin, Yang & Tokuda; *B. shianguei* Lin, Yang & Tokuda; and *B. turoguei* Lin, Yang & Tokuda in the matrix of Garcia et al. [1]. These species were studied only through literature data.

Two Australian species, *B. bursaria* (Felt) and *B. orientalis* (Felt) were tentatively transferred to *Bruggmanniella* in Kolesik and Gagné [10] based on the two separate teeth of the gonostylus. These species have unknown immature stages and one of them is only known from the male. We would rather not include them in our analysis, since an excessive amount of missing data decreases the robustness of the phylogeny.

The cladistic analysis was performed under the parsimony criterion with implicit weighting in TNT v1.5 (Willi Hennig Society Edition) [11], following the parameters described in Garcia et al. [1]. The resulting tree was displayed in Adobe Illustrator CC software (17.0). The character list follows Garcia et al. [1], with information about each character and performance values (CI, RI, *fit*) updated (Supporting Information S1 List). No character has been added or removed from the list. See Supporting Information S2 Table for normalized data, S3 Table for matrix, and S4 File for input matrix script.

## Results

The cladistic analysis under equal weighting results in 9 parsimonious trees with 172 steps. The analysis under implied weight for *k* = 3 results in a single and most stable topology (length = 179 steps, CI = 0.38, RI = 0.57, fit = 29.55,) that will be discussed here. The MPT for *k* = 3 is shown in Fig 1.

**Fig 1.**
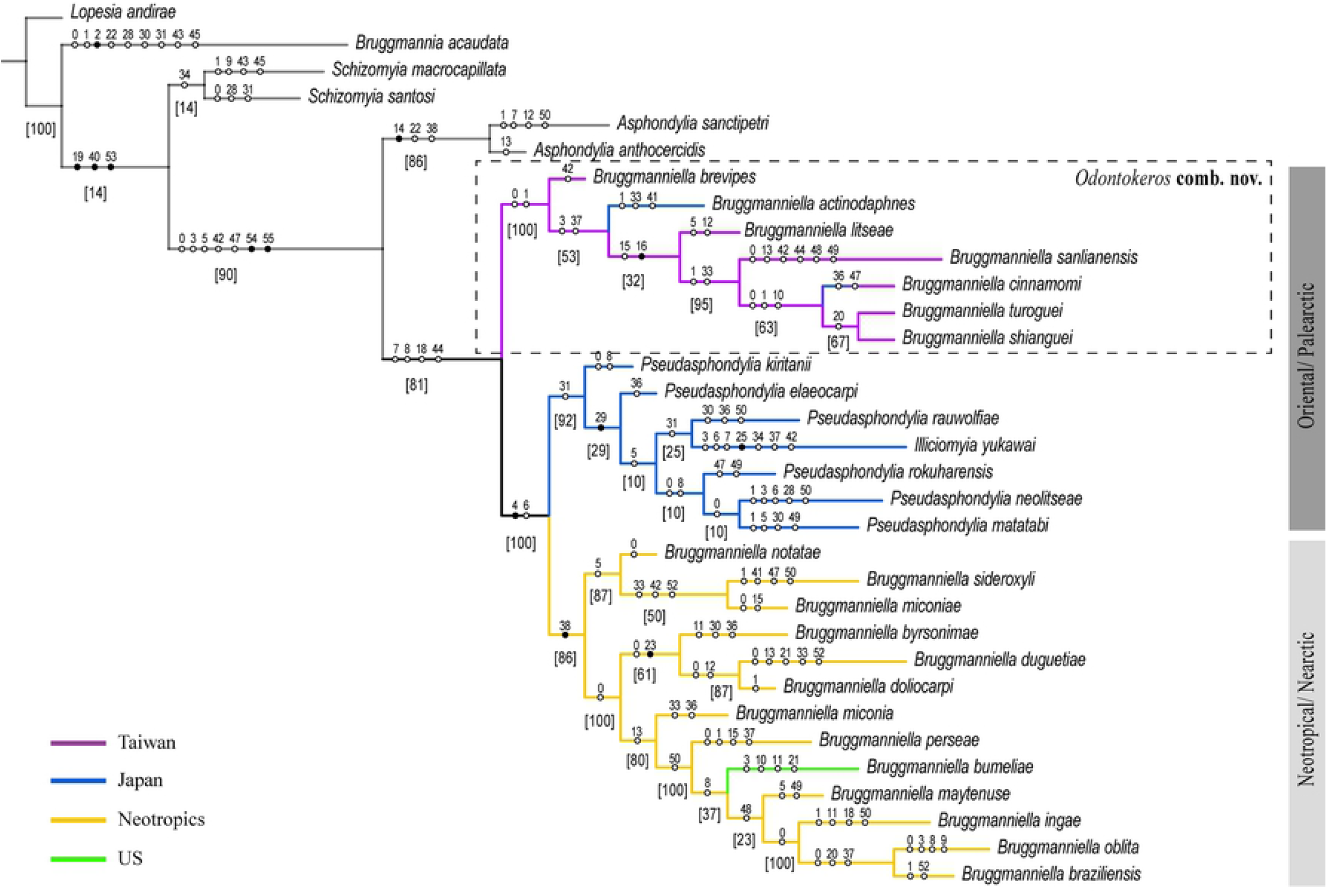
Most parsimonious tree under implied weight (*k* = 3) of the genus *Bruggmanniella* Tavares, 1909. Values in brackets show the relative Bremer support (in percentage). Purple clades symbolize the species distributed in Taiwan. Blue symbolizes the species distributed in Japan. Yellow clades represent the species with distribution in the Neotropics, mostly Brazil. Green clade represents the species with occurrence in the US.

The phylogenetic reconstruction shows the Neotropical (here including also one Nearctic species) and Asian species of *Bruggmanniella*, *Pseudasphondylia*, and *Illiciomyia* as a monophyletic group (Fig. 1) supported by four synapomorphies: one pair of inner lateral papillae and terminal papillae of the larva indistinct or absent, spiniform shape of abdominal spiracles in pupa, and gonostyli bi-toothed in males. As mentioned before, these three genera are largely recognized as a closely related group in literature [1, 2–6, 8].

The monophyly of (*Pseudasphondylia* + Neotropical *Bruggmanniella*) is congruent with the hypothesis presented in Garcia et al. [1]. The clade is strongly supported by the larval characters: outer apical teeth of spatula larger than inner ones (or inner completely reduced) and reduction of outer lateral papillae to one pair.

The *Pseudasphondylia* species are grouped by the fusion of the first and second female flagellomeres. In this analysis, *Illiciomyia* appeared nested inside *Pseudasphondylia* clade. However, more studies with other character set need to be run to further elucidate this arrangement. This monotypic genus is only distributed in Japan and induces galls on leaves, as most species of *Pseudasphondylia*, indicating that this position may not be erroneous.

## Discussion

Unlike the proposal of Garcia et al. [1], *B. actinodaphnes* and *B. cinnamomi* are not placed among the *Pseudasphondylia* species. Lin et al. [8] also suggested that *B. actinodaphnes* and *B. cinnamomi*, as all Asian *Bruggmanniella*, could not belong to *Bruggmanniella*. Our current result shows that these species do not fit under *Bruggmanniella*. It reinforces the pertinence of the genus *Odontokeros* to house not only *B. brevipes*, but all species occurring in the Oriental/Palearctic region. This clade is highly supported by the main synapomorphies: pupal antennal horn elongated, pupal cephalic setae reduced, and the molecular similarities indicated in Lin et al. [8].

Lin et al. [8] used two arguments to disqualify the analysis of Garcia et al. [1]: the branch support index and the monophyly of the Asian *Bruggmanniella* + *Pseudasphondylia* indicated by molecular sequences. The authors stated that the branches were supported by *bootstrap*, when the support used in that study was the *relative Bremer*. Although we agree that the support values of the topology of Garcia et al. [1] are low, the current analysis shows the clades of the Asian *Bruggmanniella* and (*Pseudasphondylia* + Neotropical *Bruggmanniella*) with high values of *relative Bremer index* – 100 for both clades.

Regarding the molecular proximity proposed between Asian *Bruggmanniella* and *Pseudasphondylia*, we have a different interpretation of the same results. Lin et al. [8] did not include in their analysis any Neotropical species, which leads the Asian species to be positioned as sister groups in their tree. This occurred because *Bruggmanniella*, *Pseudasphondylia*, and *Illiciomyia* are part of a group that is strongly monophyletic. The relationship among these four groups could not be tested in Lin et al. [8] analysis, as *Illiciomyia* and the Neotropical *Bruggmanniella* were not included, despite the existence of COI sequences of *Illiciomyia* and *B. miconia* available in the GenBank.

### Geographical distribution and Niche occupation

The Asian *Bruggmanniella* is mostly distributed in Taiwan (except for *B. actinodaphnes*, only found in Japan so far) and induces galls exclusively on leaves or stems in Lauraceae species, chiefly of the genus *Cinnamomum* Schaeff. We agree with Tokuda and Yukawa [4,5] and Lin et al. [8] that these Lauraceae-associated species are closely related (they occupy very similar positions in the phylogenetic trees of both studies), but they do not belong to the same genus as the Neotropical (plus Nearctic) *Bruggmanniella*.

The Asian *Bruggmanniella* occupies stems (mostly) or leaves only in species of the plant family Lauraceae, showing that the phylogenetic relationships among host species drives this association. The Neotropical *Bruggmanniella* and *Pseudasphondylia* are associated with 11 different plant families each. *Pseudasphondylia* induce galls on leaves, fruits, and flower buds while the Neotropical *Bruggmanniella* species induce gall on stems, fruits, and flower buds. The niche tissue type seems to be crucial in the association with the host plant in these two groups while phylogenetic identity of the host is less important.

The progressive decrease from four to two teeth of the spatula and in the size of the antennal horn in *Pseudasphondylia* suggest that the females lay their eggs in soft tissues (as parenchyma in leaves, fruits, and flower buds). The Neotropical *Bruggmanniella* shows an intermediate condition (the majority in stems, but some species are gall inducers in soft tissues) reflected by the decrease from four to three teeth of the spatula in some species (the type-species *B. braziliensis* and *B. oblita*) and the less developed antennal horns.

*Bruggmanniella* is confirmed here as a monophyletic and divergent group from the Asian species, based on the loss of the parameres. This result also allows us to understand the presence of parameres as a plesiomorphic state to the Asphondyliina subtribe. The heterogeneity of ecological niches and the disjunct distribution of the Asian and Neotropical species of *Bruggmanniella* were better understood. A more comprehensive study including all species of Asphondyliina with two-toothed gonostyli, using a set of different molecular markers concatenated with morphological characters, will greatly contribute to understanding this interesting evolutionary history.

### Taxonomy

For all the arguments presented here, we re-established the combination of *Odontokeros brevipes* comb. rev., and transfer the species *B. actinodaphnes, B. cinnamomi, B. brevipes, B. litseae, B. sanlianensis, B. shianguei*, and *B. turoguei* to the same genus: *O. actinodaphnes* **comb. nov.**, *O. cinnamomi* **comb. nov.**, *O. brevipes* **comb. nov.**, *O. litseae* **comb. nov.**, *O. sanlianensis* **comb. nov.**, *O. shianguei* **comb. nov.**, and *O. turoguei* **comb. nov.**

*Odontokeros* – Diagnosis. Prothoracic spatula with 2 or 4-teeth, if four, **inner teeth shorter than outer ones**; pupa with deeply toothed antennal horns and prothoracic spiracles well developed, upper pupal antennal horn elongated and pupal cephalic setae reduced, **parameres membranous**, and molecular similarities indicated in Lin et al. [8].

*Pseudasphondylia* – Prothoracic spatula with 2 or 4-teeth, if four, **inner teeth larger than outer ones**; first and second female flagellomeres fused, **parameres membranous**, bi-toothed spatula, if four, inner teeth shorter than outer.

*Bruggmanniela* – Prothoracic larval spatula with 3 or 4-teeth, inner teeth (or tooth) **larger than outer ones**; pupa with antennal horns and well-developed prothoracic spiracles, upper and frontal horns absent, pupal cephalic margin thickened; male genitalia with two-toothed gonostyli, **parameres absent**; cerci-like lobes on female abdominal segment VIII.

## Acknowledgment

The authors acknowledge Dr. John Wenzel for helpful comments on the draft version of the manuscript.

## Supporting Information

**S1 List – Characters**

**S2 Table – Normalized Data**

**S3 Table – Matrix**

**S4 File – Input Matrix Script**

